# METCAM/MUC18 plays a tumor promoter role in the development of nasopharyngeal carcinoma type III

**DOI:** 10.1101/2024.12.07.627349

**Authors:** Yen-Chun Liu, Chun-Chuan Ko, Yu-Jen Chen, Guang-Jer Wu

## Abstract

Our previous studies by testing the effects of huMETCAM/MUC18 over-expression on the tumorigenesis of the NPC-TW01 cell line supports the notion that huMETCAM/MUC18 plays a tumor suppressor role in the development of nasopharyngeal carcinoma (NPC) type I. However, it is not clear if this notion also holds true for another NPC cell line, NPC-TW04, which was established from NPC type III. For this purpose, we further investigated the effects of huMETCAM/MUC18 over-expression in the NPC-TW04 cell line on their *in vitro* cellular behaviors and *in vivo* tumorigenesis in athymic nude mice. First, the G418-resistant clones were obtained after transfection of the huMETCAM/MUC18 cDNA into the NPC-TW04 cell line. Then, one NPC-TW04 clone, which expressed a high level of huMETCAM/MUC18, and one empty vector (control) clone were used for testing effects of huMETCAM/MUC18 over-expression on their *in vitro* behaviors (intrinsic growth rate, motility, and invasiveness, and growth on three-dimensional basement membrane) and *in vivo* tumorigenesis (via subcutaneous injection) in the athymic nude mice. Time course of tumor proliferation and final tumor volumes/weights were determined. Tumor sections were used for histology and immunohistochemistry (IHC) analyses. Tumor lysates were used to determine levels of huMETCAM/MUC18 and various downstream key effectors. We found that over-expression of huMETCAM/MUC18 increased *in vitro* motility and invasiveness, induced the invasive growth in the 3D basement membrane culture assay, and increased *in vivo* tumorigenicity of NPC-TW04 cell line. The expression levels of downstream effectors, such as an anti-apoptosis index, proliferation index, an index of metabolic switch to aerobic glycolysis, angiogenesis indexes and survival pathway index, were increased, whereas the pro-apoptosis index decreased in the tumors. Thus, contrary to the role of huMETCAM/MUC18 in the NPC-TW01 cells, huMETCAM/MUC18 over-expression in the NPC-TW04 cells increased epithelial-to-mesenchymal transition and also *in vitro* invasive growth and *in vivo* tumorigenesis, suggesting that it plays a tumor promoter role in the development of NPC type III by reducing apoptosis and increasing anti-apoptosis, angiogenesis, proliferation and metabolic switch to aerobic glycolysis.

## 1. INTRODUCTION

Nasopharyngeal carcinoma (NPC) is a non-lymphomatous, squamous-cell carcinoma that usually develops around the ostium of the Eustachian tube in the lateral wall of the nasopharynx [1]. NPC is composed of phenotypically and genotypically heterogeneous populations of neoplastic cells; thus, it usually manifests one of the three histological subtypes: keratinizing squamous cell carcinomas (WHO type I), nonkeratinizing squamous cell carcinomas (WHO type II), and undifferentiated carcinomas (WHO type III) [2]. Nonkeratinizing NPC is the most common subtype, accounting for >95% of cases in endemic areas [2].

NPC is a rare head and neck cancer throughout most of the world but common in certain geographic areas, such as Southeast Asia, China (Guangdong province), Taiwan, Hong Kong, Northeastern region of India, the Middle East, North Africa, Inuit of Alaska, and native Greenlanders [1, 3, 4]. NPC affects both the young and the elderly patients and has caused a significant burden on cancer statistics [5, 6]. Though rare among whites, NPC is particularly common in Chinese emigrants continue to have a high incidence of the disease, but the rate of NPC among ethnic Chinese born in North America is considerably lower than those born in China [1, 3–6]. Types II and III NPC are Epstein–Barr virus (EBV) associated and have better prognoses than type I; EBV infection is generally absent in type I, especially in nonendemic areas [1, 3–6]. Thus, epidemiologic risk factors important for the development of NPC include genetic susceptibility and life styles/environmental factors. Genetic susceptibility includes genetic polymorphisms in certain human leukocyte antigen subtype, such as HLA subtypes. The environmental factors include exposure to nitrosamines in salted and pickled foods. Furthermore, the Epstein–Barr virus in particular has been implicated in the molecular abnormalities leading to NPC, since almost all NPC tumors, regardless of histologic subtype, have comorbid EBV infections [1, 3–6]. However, the exact mechanisms remain elusive; the tumorigenesis of NPC may be due to the consequence of a complex interplay among these risk factors and is resulting from a sequential multistep process that transforms normal nasopharyngeal epithelium into cancer cells [1, 3–6]. The key events include carcinogen exposure, mutations in genes regulating immune responses, epigenetic processes, and signal transduction, as well as the genome instability promoted by the EBV [1, 4–6].

NPC patients at early stages can be effectively treated by radiotherapy and concurrent adjuvant chemotherapy [7–8]. High survival rates are reported for stage 1 and 2 diseases; however, most NPC patients are diagnosed at late stages, at when the curative rate of various treatments is low with a disease relapse rate as high as 82% [7–8]. The main reasons for the majority of NPC being diagnosed at an advanced stage are due to non-specific presenting symptoms (cervical nodal enlargement, headache, nasal and aural dysfunction), delay in seeking treatment after the onset of symptoms, and the difficulty of a thorough nasopharyngeal examination [4–8]. Thus, more targeted treatments of NPC are necessary to be developed. However, this is impossible without a specific biomarker for each type of NPC; thus, the diagnosis of NPC patients at early stage is impossible or the prediction of relapse is often too late [9]. From our previous studies, we suggest that METCAM/MUC18 may fulfill this need [10–11].

Human METCAM/MUC18 (huMETCAM/MUC18), a cell adhesion molecule (CAM) in the immunoglobulin gene super-family [12–13], is expressed in the normal nasopharynx [10], as well as in several other normal tissues [14–16]. From our previous studies, huMETCAM/MUC18 plays an interesting dual role in malignant progression of many epithelial tumors [17]: as a tumor and metastasis promoter in breast cancer [15, 18–20] and prostate cancer [21–28] and a metastasis promoter in melanoma [29–31], but as a tumor and metastasis suppressor in ovarian cancer [32–35] and one mouse melanoma cell line [36]. Nevertheless, it is not known if huMETCAM/MUC18 plays any role in the progression of nasopharyngeal carcinoma (NPC).

For this purpose, we first initiated the immunohistochemical studies of normal nasopharynx and three types pf NPC and found that the reduced expression of huMETCAM/MUC18 correlates with the emergence of all the three subtypes of NPC, suggesting the notion that huMETCAM/MUC18 may play a negative role in the progression of the cancer [10]. We then tested the effects of over-expression of huMETCAM/MUC18 on *in vitro* cellular behaviors and *in vivo* tumorigenesis of NPC-TW01 cells, which were established from NPC Type I, the results support the notion that huMETCAM/MUC18 indeed plays a tumor suppressor role in the development of NPC Type I [11], consistent with the above negative correlation [10]. However, we did not know if this notion is also generally applicable to another cell line, NPC-TW04, which was established from NPC type III [37, 38].

In this report we further scrutinize the above notion by expanding our investigation to test the effects of huMETCAM/MUC18 over-expression on *in vitro* cellular behaviors and *in vivo* tumorigenesis of another NPC cell line, NPC-TW04, in athymic nude mice. In contrast to the results obtained by using the NPC-TW01 cell line [11], we obtained both *in vitro* and *in vivo* evidence to surprisingly support the opposite notion that METCAM/MUC18 over-expression promoted the tumorigenesis of NPC-TW04 cell line, hence enhanced the tumor development of the NPC type III. From the results of further preliminary mechanistical studies, we also suggest that huMETCAM/MUC18 plays a tumor promoter role in the development of NPC type III by reducing apoptosis and increasing anti-apoptosis, angiogenesis, survival, proliferation and metabolic switch to aerobic glycolysis.

## 2. MATERIALS AND METHODS

### 2.1. Cell lines and antibodies

The NPC-TW04 cell line was obtained from Dr. Chin-Tarng Lin, Department of Pathology, National Taiwan University, Taipei, Taiwan [37, 38]. The NPC-TW04 cell line was established from NPC type III and thoroughly karyotyped by him [33, 34]. The anti-huMETCAM/MUC18 antibody was prepared from chicken, which were immunized with the middle fragment of a recombinant huMETCAM/MUC18 protein overly expressed in *E. coli* [13]. This antibody could recognize the epitopes of the huMETCAM/MUC18 protein expressed in many human cancer cell lines and in formalin-fixed, paraffinized tissue sections with a minimal interference from glycans because this portion of the huMETCAM/MUC18 protein lacks any N-glycosylation sites [13]. Furthermore, this antibody has a high specificity with a minimal cross-reactivity with the mouse METCAM/MUC18 protein [39].

### 2.2. Growth of various human cancer cell lines

The positive control, the human melanoma cell line, SK-Mel-28 from ATCC, was maintained in Eagle’s MEM supplemented with sodium pyruvate, additional non-essential amino acids and vitamins, and 10% fetal bovine serum and with penicillin and streptomycin and in 5% CO_2_ in a 37° C incubator. The NPC-TW04 cell line was maintained in DMEM (4.5% glucose) supplemented with 5% fetal bovine serum and 100 μg/ml kanamycin in 5% CO_2_ and in a 37° C incubator [37, 38]. All the G418-resistant (G418^R^) clones derived from NPC-TW04 were grown in the same medium plus 0.4 mg/ml of G418 (Geneticin, GIBCO/Life Technology). All media were from Invitrogen/Life technology and FBS from GIBCO/ Life technology or Sigma Chemical Co.

### 2.3. Lipofection of NPC-TW04 Cells and selection for METCAM/MUC18-expressing G418^R^-Clones

Lipofection was carried out with Lipofectamine 2000 (1 mg/ml, Cat #11668-019, lot#1024993, Invitrogen) according to the company-suggested procedure with minor modifications, as described [11]. Similar to NPC-TW01, G418-resistant (G418^R^) clones were transferred and expanded sequentially from 24-well to 6-well culture plates [11]. Cell lysate of each clone was made from each well of 6-well plates with the addition of Western blot lysis buffer [40]. The METCAM/MUC18-positive clones were further expanded, processed, and frozen in liquid nitrogen for preservation as stocks.

### 2.4. Determination of *in vitro* (intrinsic) growth rate of NPC-TW04 clones

The growth rates of a high-expressing G418^R^ clone and a vector-control clone were determined by direct counting of cell numbers of monolayers after treatment with trypsin, as described [11]. The averages of the hex plicate cell numbers from each time point were used for calculation of the growth rate. The relative growth rate was determined between three sets of two-time points [11].

### 2.5. Cell motility assay

A cell motility assay was carried out according to a published method [41] with minor modifications [11] by using the Boyden type Trans-well system (Becton/Dickinson Falcon 35-3503). The average of three repeats is shown.

### 2.6. Cell invasiveness assay

A cell invasiveness assay was carried out according to a published method [41] with minor modifications [11] by using the Boyden type Trans-well system (Becton/Dickinson Falcon 35-3503). The average of three repeats is shown.

### 2.7. Three-dimensional (3D) basement membrane (Matrigel) culture assay

The published procedures of a 3D embedded basement membrane culture assay [42] were followed with slight modifications as described [11].

### 2.8. Determination of *in vivo* tumorigenicity of NPC-TW04 clones/cells in athymic nude mice

The guidelines of National Institutes of Health, USA for the care and use of laboratory animals were strictly followed for all animal experiments, which were reviewed by the Mackay Memorial Hospital Experimental Animal Care and Use Committee with an approval # MMH-A-S-101-31 and CYCU Experimental Animal Care Committee with an approval #10119, as described [11]. Five 37 days old female or male athymic nude mice Balb/cAnN.Cg-Foxnlnu/Cr1Nar1 from National Laboratory Animal Center, Taipei, Taiwan, were used for the subcutaneous injection of cells from each clone. A single cell suspension from monolayer cultures of NPC-TW04 cells of the clone #76 (p59), or the vector clone V5 (p62) were prepared by treatment with trypsin, washed with the phosphate-buffered saline, and counted by a hemocytometer. 5 x 10^5^ cells were re-suspended in 0.05 ml of cold DMEM medium without fetal bovine serum, mixed with an equal volume of 15.44-15.95 mg/ml of Cutrex [11, 42, 43], and used for injection to each mouse. Mice were anesthetized *IP* with Zoletil 50 (50 mg/ mouse) and subcutaneously injected with the cells at the back of the neck region or right leg femur by using gauge #30G_1/2_ needles. After injection, the well-being of the mice was daily checked, and the size of tumor was weekly measured with a caliper till the endpoint of the experiment (40 days). There was no adverse effect for the mice bearing tumors till the endpoint. Tumor volume was calculated by using the ellipsoid formula V = π/6 (d1 x d2)^3/2^ (mm^3^) [41]. At the endpoint, all the mice were euthanatized with 100% carbon dioxide in an inhalation chamber and the tumor from each mouse was excised, weighed, and a major portion of it was used for making cell lysate for Western blot analysis. A small portion of the tumor was fixed in 10% formaldehyde (Cat #A3684, 2500, Lot# 2T001138, phosphate-buffered histology grade, AppliChem GmbH, Germany), paraffinized, and sectioned for histology (Leica Microsystems) and immunohistochemistry.

### 2.9. Western blot analysis

Cell extracts from various cultured cell lines were prepared by directly lysing the monolayer cells with the Western blot lysis buffer, as previously described [11, 40]. The Western blot lysate from each NPC xenograft tumor tissue was prepared from the homogenate of the tissue, as also previously described [11]. HuMETCAM/MUC18 protein expression in cellular and tumor tissue extracts (10 μg/lane) was determined by the standard procedure of Western blot analysis by using our chicken anti-huMETCAM/MUC18 protein IgY as the primary antibody (1/300 dilution of 15 mg/ml) and the AP-conjugated rabbit anti-chicken IgY (AP162A, Chemicon) as the secondary antibody (1/2000 dilution of 2 mg/ml) [11]. The same membrane was subsequently used for the detection of the expression of the house-keeping genes, actin and β-tubulin, as the loading controls [11]. The primary antibodies and AP-conjugated secondary antibodies to detect expression levels of Bcl2, Bax, PCNA, VEGF, pan-AKT, phospho-AKT (SER437), and LDH-A were previously described [11]. The substrates BCIP/NBT (S3771, Promega) were used for the color development. The image of the huMETCAM/MUC18 band and all other bands on membranes were scanned with an Epson Photo Scanner model 1260 and their intensities were quantitatively determined by a NIH Image J program version 1.31.

### 2.10. Histology and Immunohistochemistry

Paraffin-embedded tissue sections (5 μm) were de-paraffinized, rehydrated with graded alcohol and PBS, and used for histological staining and immunohistochemical analyses, as previously described [11]. A tissue section of the subcutaneous tumor derived from the LNCaP-expressing clone, LNS239, [26] was used as a positive external control for the immunohistochemical staining. Negative controls had the primary antibody replaced by the non-fat milk or the control isotype chicken IgY.

### 2.11. Determination of vascular density in tumor sections

Vascular density in sections of SC tumors after H&E Staining were counted quantitatively under 12 microscope fields and the average number of vasculatures per field was indicated [11].

### 2.12. Statistical analysis

The student’s *t* test was used for standard deviations and One-Way ANOVA was used to analyze the statistical significance of the data in all figures. Two corresponding sets of data were considered significantly different if the *p* value was < 0.05.

## 3. RESULTS

### 3.1. Expression of huMETCAM/MUC18 protein in the G418^R^-clones of the NPC-TW04 cell line

Since NPC-TW04 cells weakly express METCAM/MUC18 in comparison to that of SK-Mel-28 cells [10, 44], we biochemically increased the expression of METCAM/MUC18 by transfecting the cells with the huMETCAM/MUC18 cDNA *via* a lipofection reagent and isolated clones that were resistant to G418 [44]. G418-resistant clones (G418^R^-clones), which expressed different levels of the protein were used for testing the hypothesis. Similar to the NPC-TW01 cell line [44], we found that Lipofectamine 2000 was more efficient than the other three lipofection reagents to successfully yield a few high-expressing, some intermediate-expressing, and some low-expressing clones [44]. Figure 1 shows the expression of METCAM/MUC18 in two typical G418^R^-clones (#76 and #72) and one vector control clone (V5 clone). The clones #76 and #72 expressed 62% and 23% of the protein, respectively, whereas a vector control clone (V5) expressed about 9% of the protein, assuming that the SK-Mel-28 cell line expressed 100% of the protein.

**Figure 1.**
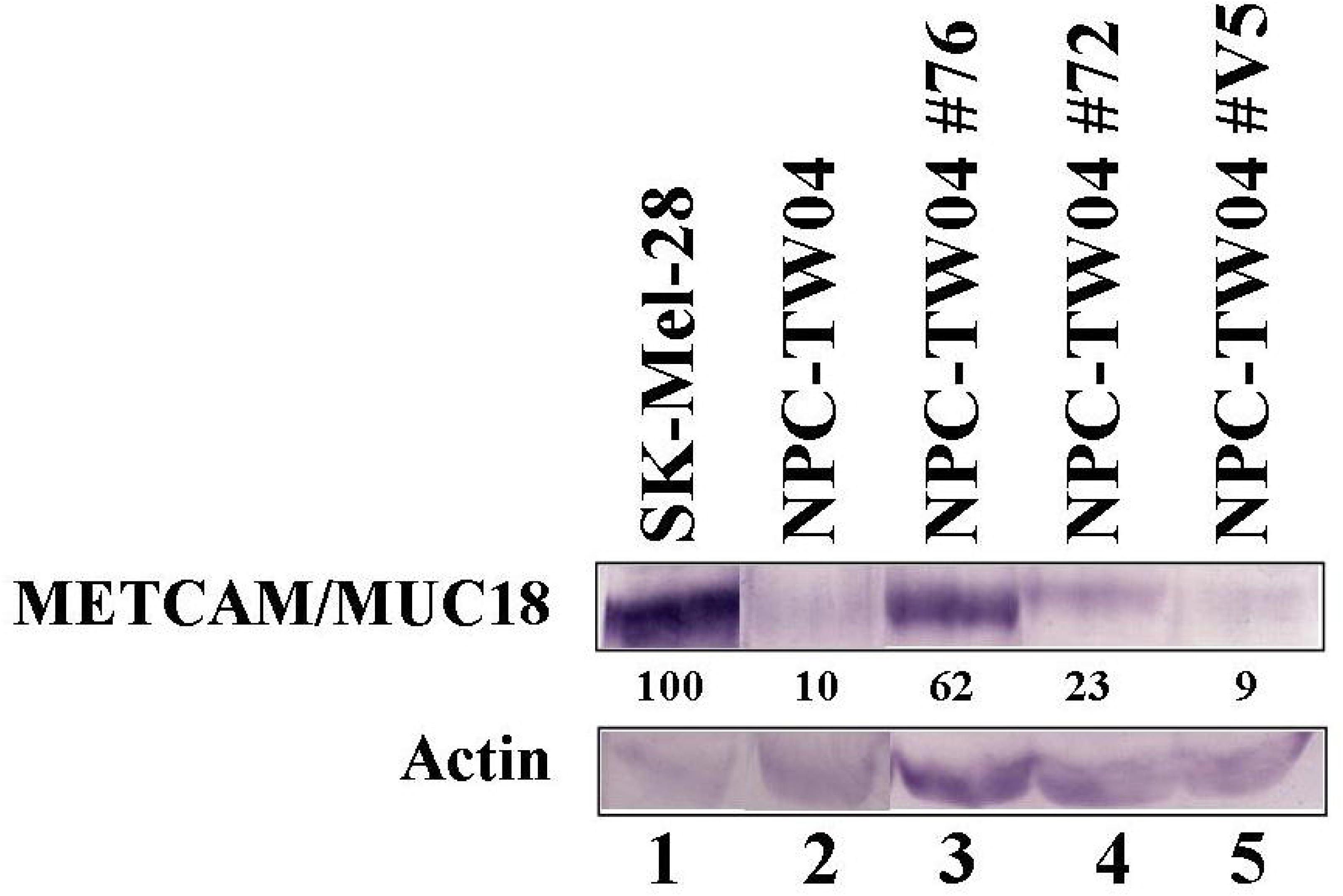
Expression of the huMETCAM/MUC18 protein in the NPC-TW04 G418^R^ clones. Human METCAM/MUC18 expression in lysates prepared from various clones/cells was determined by Western blot analysis as described in the Section 2. The METCAM/MUC18 expression level in the cell lysate from a human melanoma cell line, SK-Mel-28, was used as a positive control (lane 1), and that from the parental human nasopharyngeal cancer cell line, NPC-TW04, was used as a negative control (lane 2). The METCAM/MUC18 expression in cell lysates from two METCAM/MUC18 cDNA-transfected NPC-TW04 clones (METCAM Clones #76 and #72) and one vector control clone V5 are shown in lanes 3-5. The METCAM clones #76 and #72 were de-rived from NPC-TW04 cells transfected with the human METCAM/MUC18 cDNA gene. The vector control clone V5 was from NPC-TW04 cells transfected with the empty vector. Actin is shown as the loading control. The number under each lane indicates the relative level of METCAM/MUC18 of each cell line after correction for the loading control of the house-keeping gene, actin, assuming that of the SK-Mel-28 was 100%.

### 3.2. Over-expression of huMETCAM/MUC18 did not affect the *in vitro* growth rates of NPC-TW04 clones/cells

We tested whether the over-expression of huMETCAM/MUC18 affected the growth of NPC-TW04 cells *in vitro*. As shown in Figure 2, the over-expression of huMETCAM/MUC18 did not significantly affect the growth rate of the cells, since the growth rate of the clone #76 was statistically not much different from that of the vector control clone V5. We thus concluded that the over-expression of huMETCAM/MUC18 in the NPC-TW04 cells did not change the *in vitro* intrinsic growth rate.

**Figure 2.**
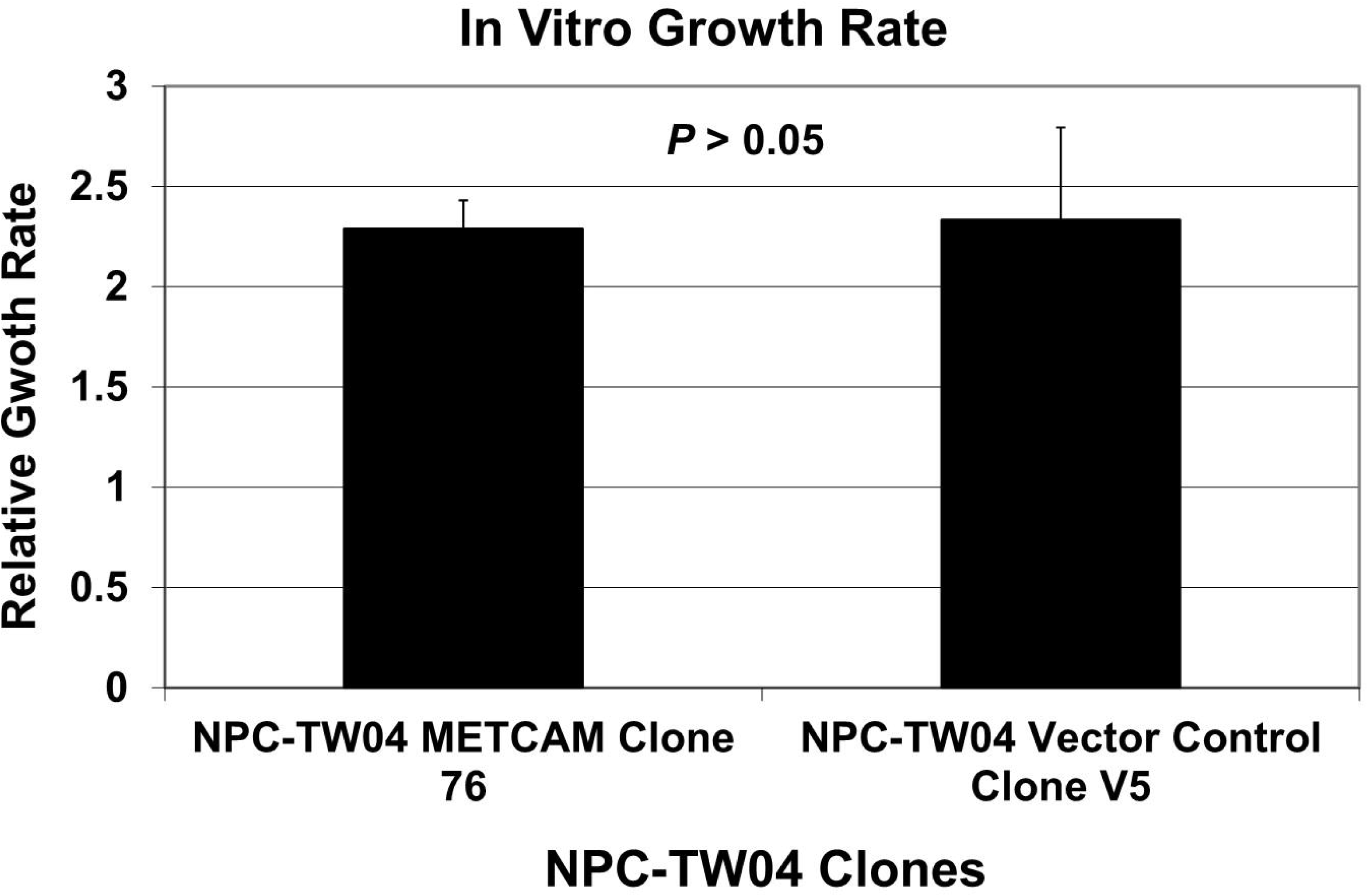
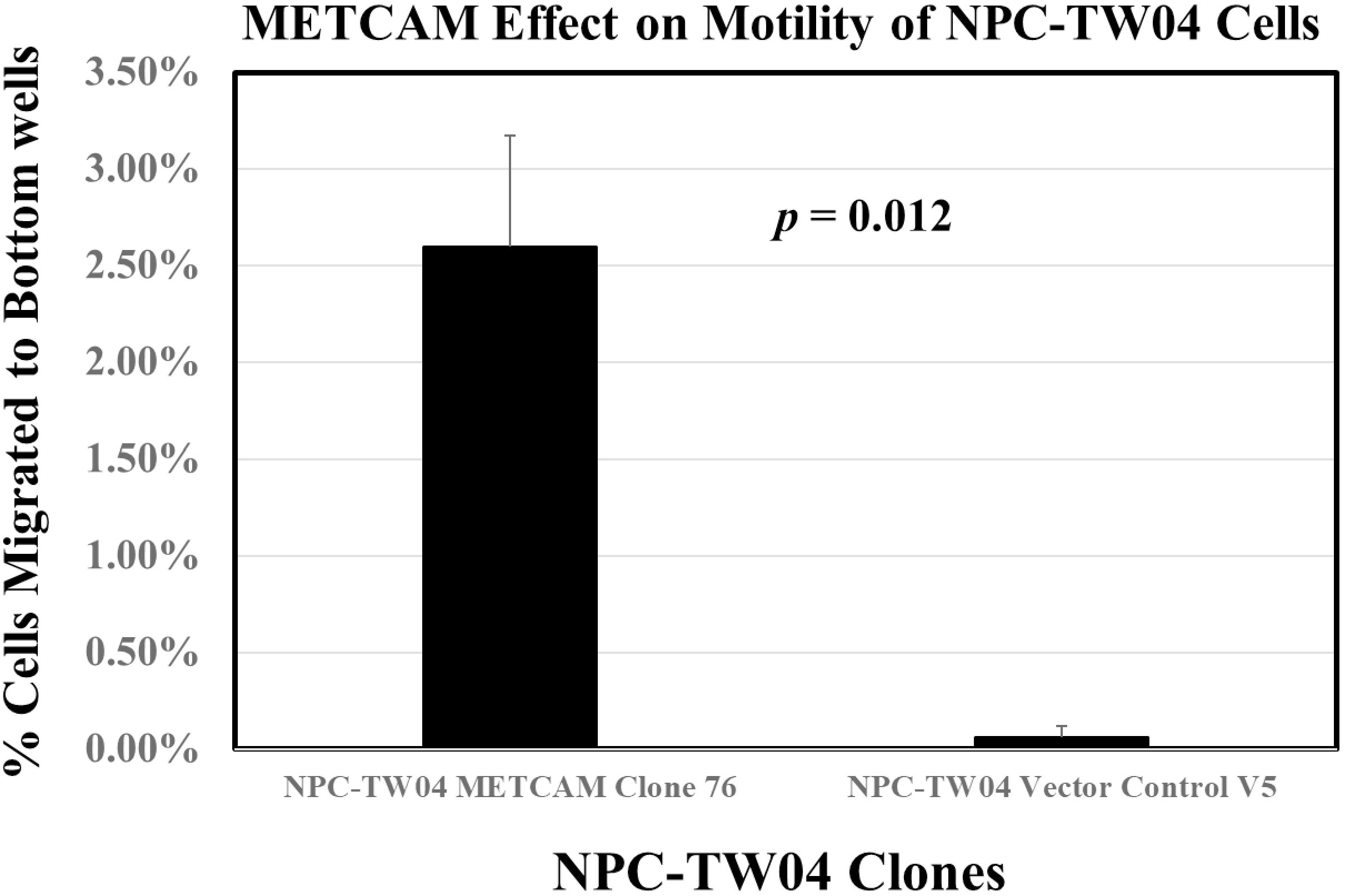
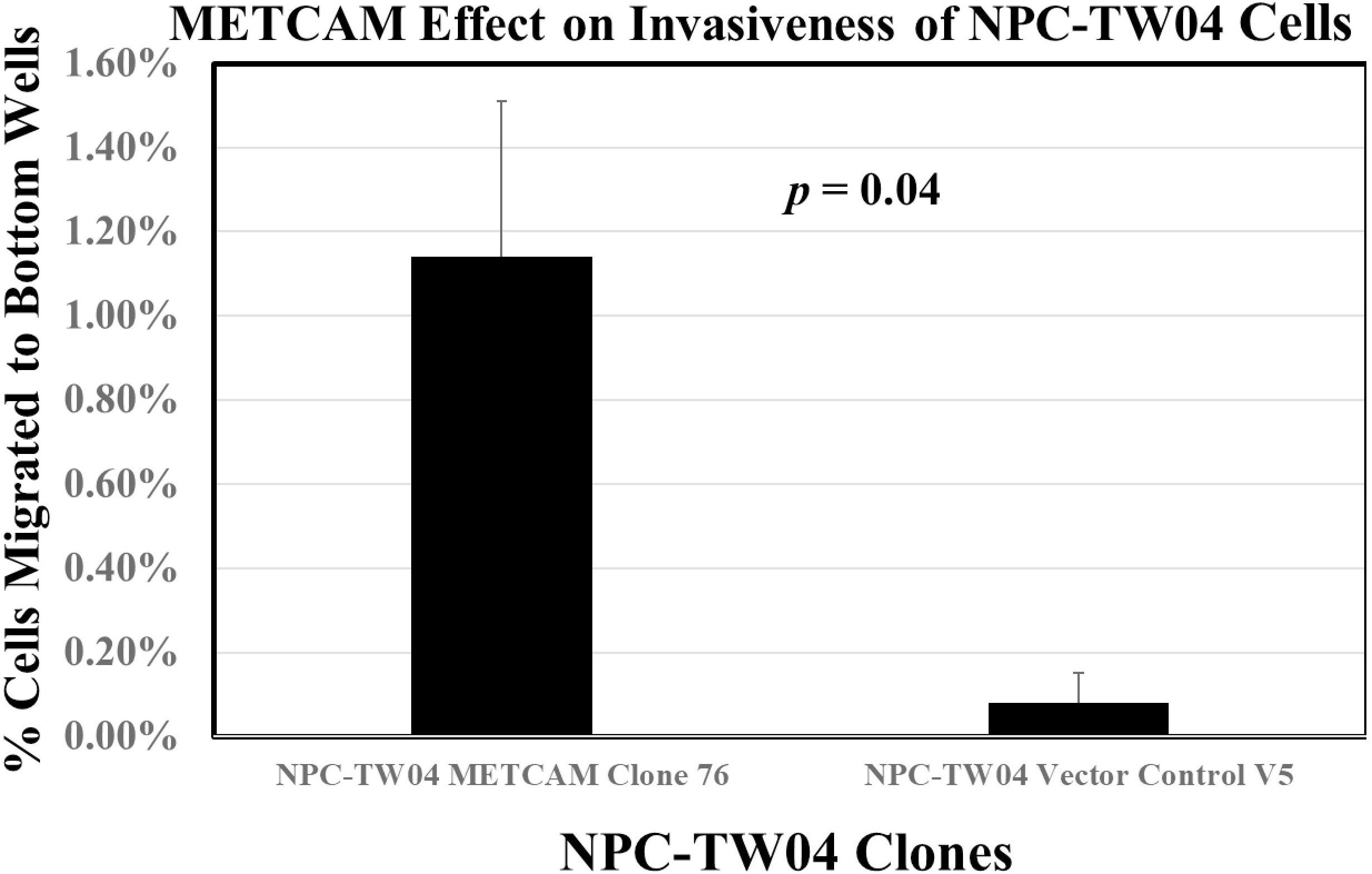
*In vitro* growth rates of NPC-TW04 clones/cells. Effects of huMETCAM/MUC18 expression on the *in vitro* growth rate of NPC-TW04 cells were carried out by determining the growth rate of the NPC-TW01 clones 76 and V5 by direct counting the cells at 24, 48, 72, and 96 hours after seeding the cells as described in the Section 2. The mean and standard deviation of three relative growth rates for each clone was plotted. The *p* value of the results was >0.05, indicating the difference is insignificant.

### 3.3. Overexpression of huMETCAM/MUC18 increases the *in vitro* motility and invasiveness of NPC-TW04 cells

Figure 3A shows that the high huMETCAM/MUC18-expressing clone (#76) had a 44-fold higher motility than the vector control clone (V5). Thus, over-expression of huMETCAM/MUC18 significantly increased the motility of NPC-TW04 cells. Figure 3B shows that the high huMETCAM/MUC18-expressing clone (#76) had a 10-fold higher invasiveness than the vector control clone. Thus, over-expression of huMETCAM/MUC18 also significantly increased the invasiveness of NPC-TW04 cells. Taken together, we concluded that over-expression of huMETCAM/MUC18 augmented both *in vitro* motility and *in vitro* invasiveness of NPC-TW04 cells.

**Figure 3.**
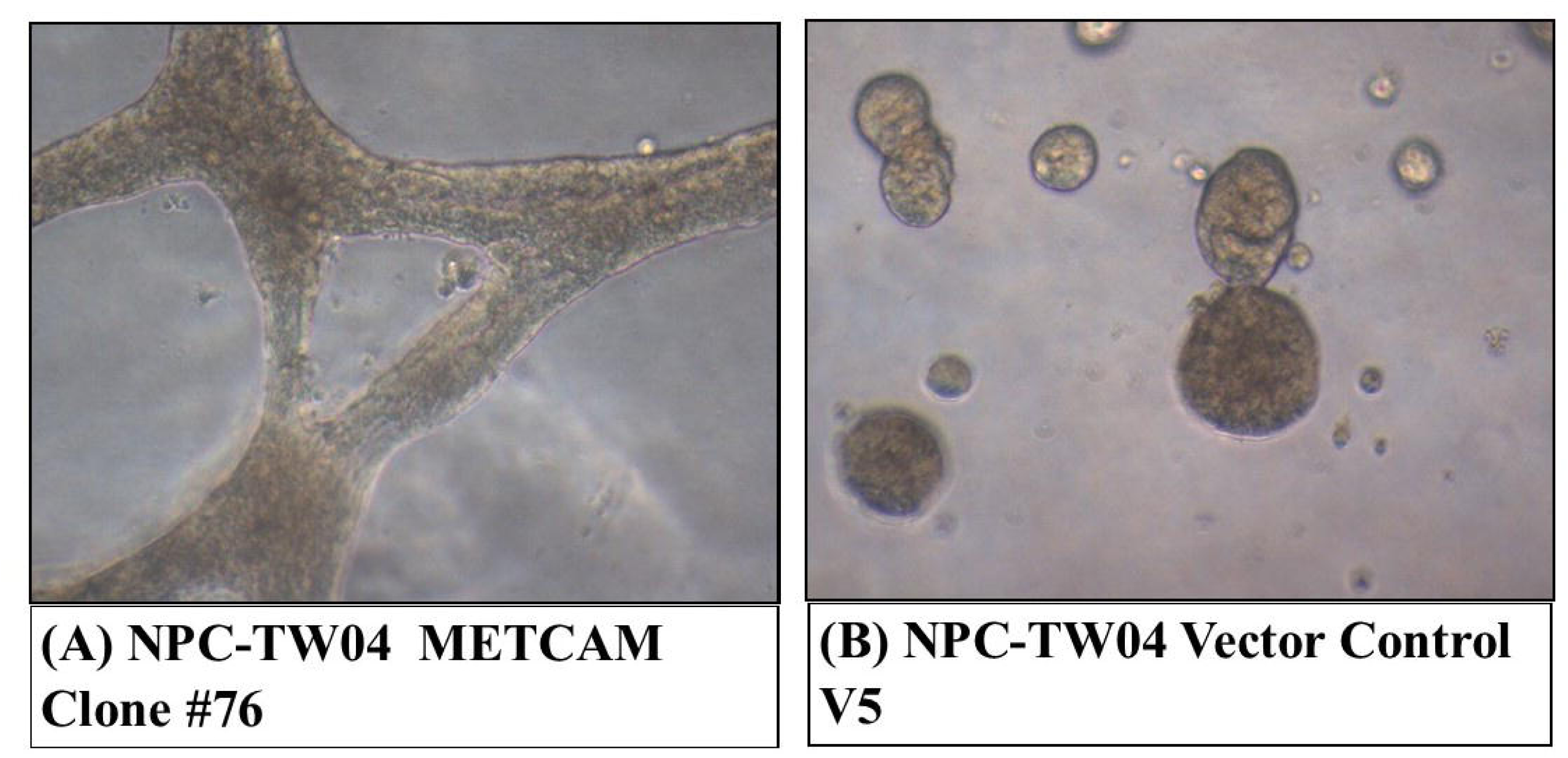
Effects of huMETCAM/MUC18 over-expression on the *in vitro* motility (A) and invasiveness (B) of NPC-TW04 cells. All *in vitro* motility and invasiveness tests were performed as described in the Section 2. The experiments were repeated three times and means and standard deviations of triplicate values are indicated. (A) Effects of huMETCAM/MUC18 expression on the *in vitro* motility of the NPC-TW04 METCAM clone #76 and the vector control clone V5 were determined. Means and standard deviations of triplicate values of the motility tests are indicated. *P* values were obtained by comparing the motility of the clone #76 with the control V5 clone. (B) Effects of huMETCAM/MUC18 expression on the *in vitro* invasiveness of the NPC-TW04 METCAM clone #76 and the vector control clone V5 were determined. Means and standard deviations of triplicate values of the invasiveness tests are indicated. *P* values were obtained by comparing the invasiveness of the clone #76 with the control V5 clone.

### 3.4. Overexpression of huMETCAM/MUC18 increases the invasive growth of the NPC-TW04 cells in the 3D basement membrane assay

Similar to the NPC-TW01 cells/clones, we were unable to demonstrate *in vitro* tumorigenesis of the cells in an *in vitro* colony formation assay in soft agar [45]. Alternatively, the 3D basement membrane assay [42] was useful not only for a similar test, but also for the growth behavior/organization of tumor cells in the 3D culture (Figure 4). Figure 4A shows that NPC-TW04 METCAM clone 76 formed an invasive growth (an extensive tubular-like growth) in the 3D basement membrane assay. In contrast, the vector control V5 formed a less aggressive growth (a spherical growth), as shown in Figure 4B. Taken together, we concluded that over-expression of huMETCAM/MUC18 appeared to induce a more invasive (aggressive) growth of NPC-TW04 cells in the 3D basement membrane culture assay.

**Figure 4.**
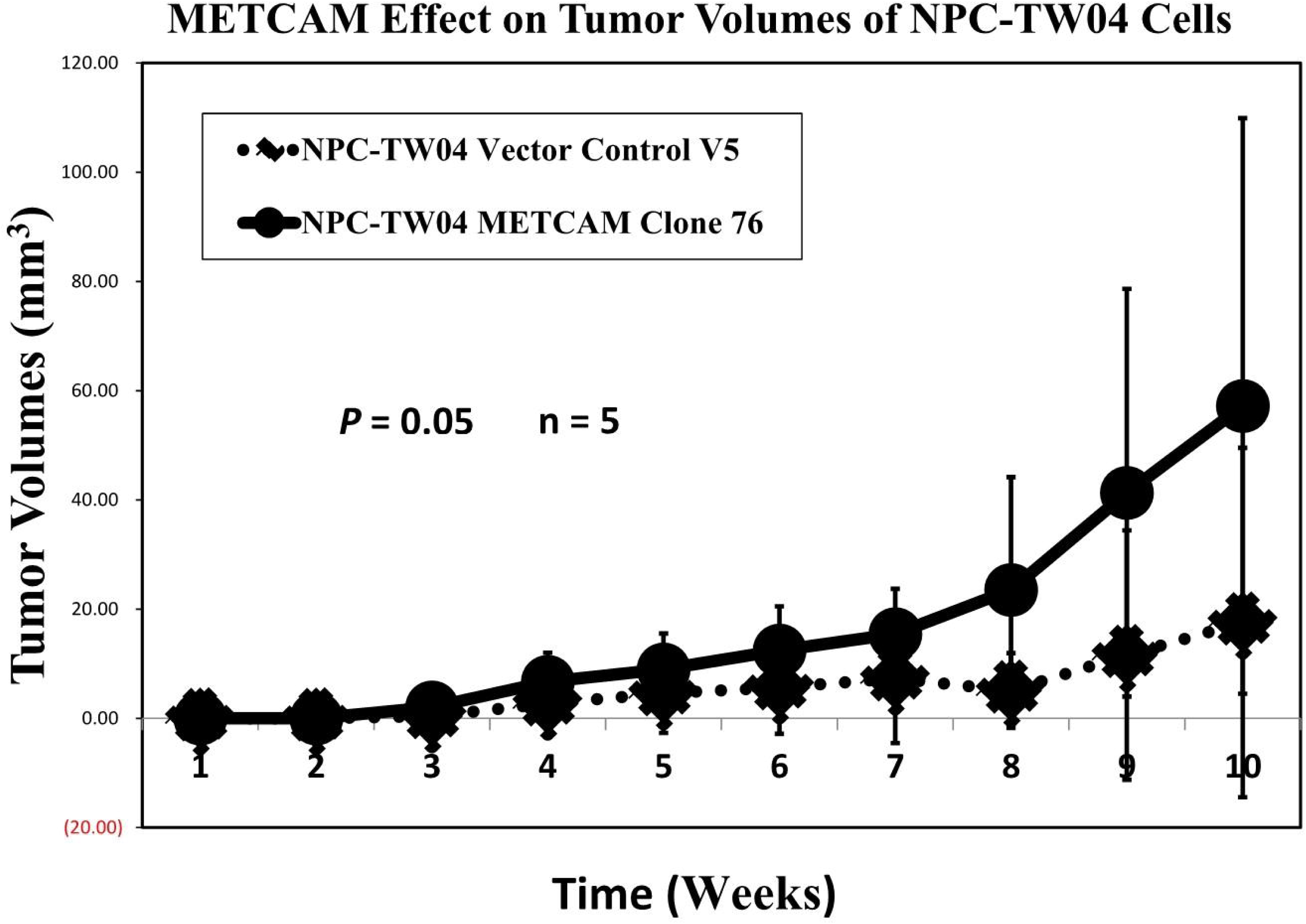
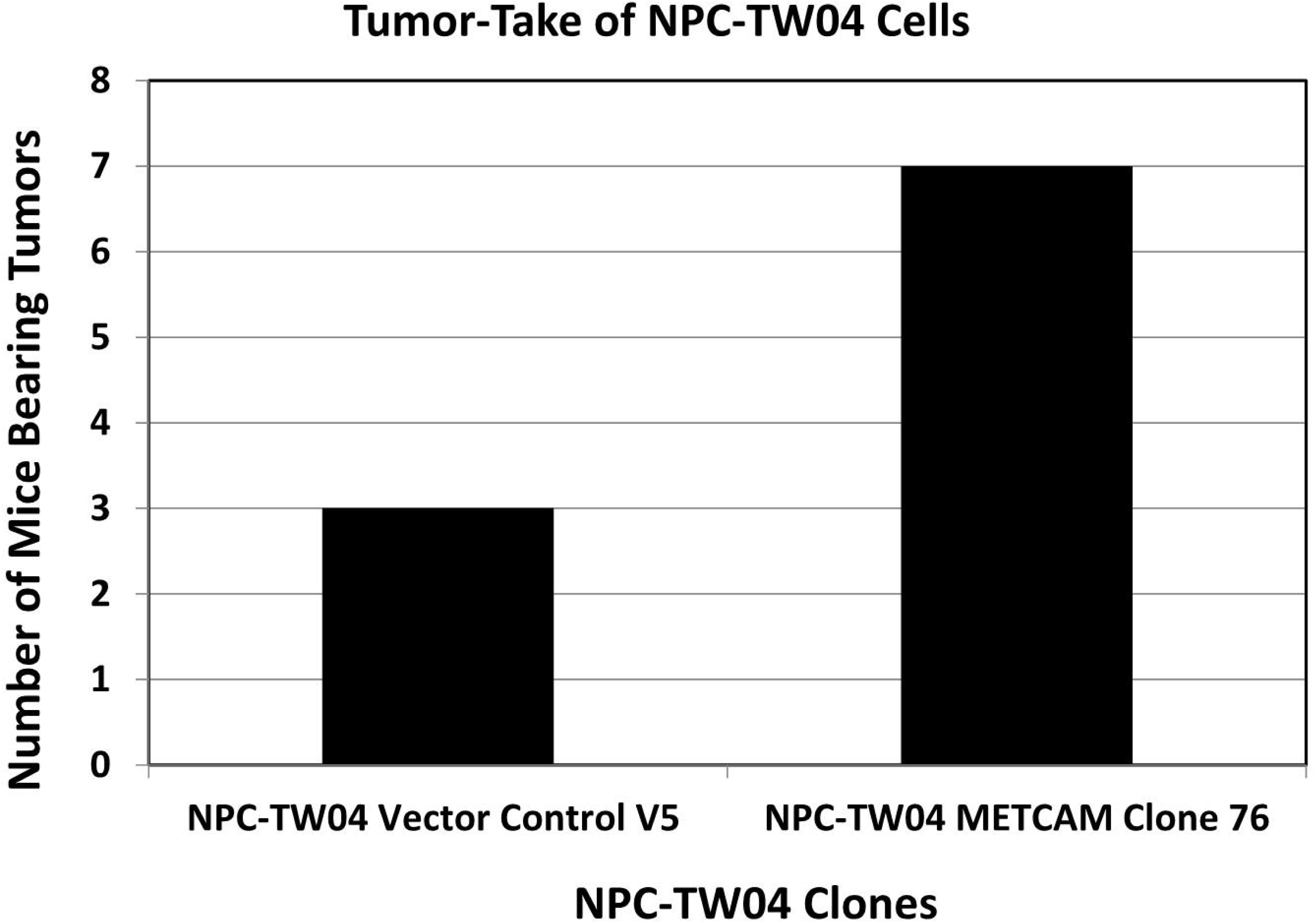
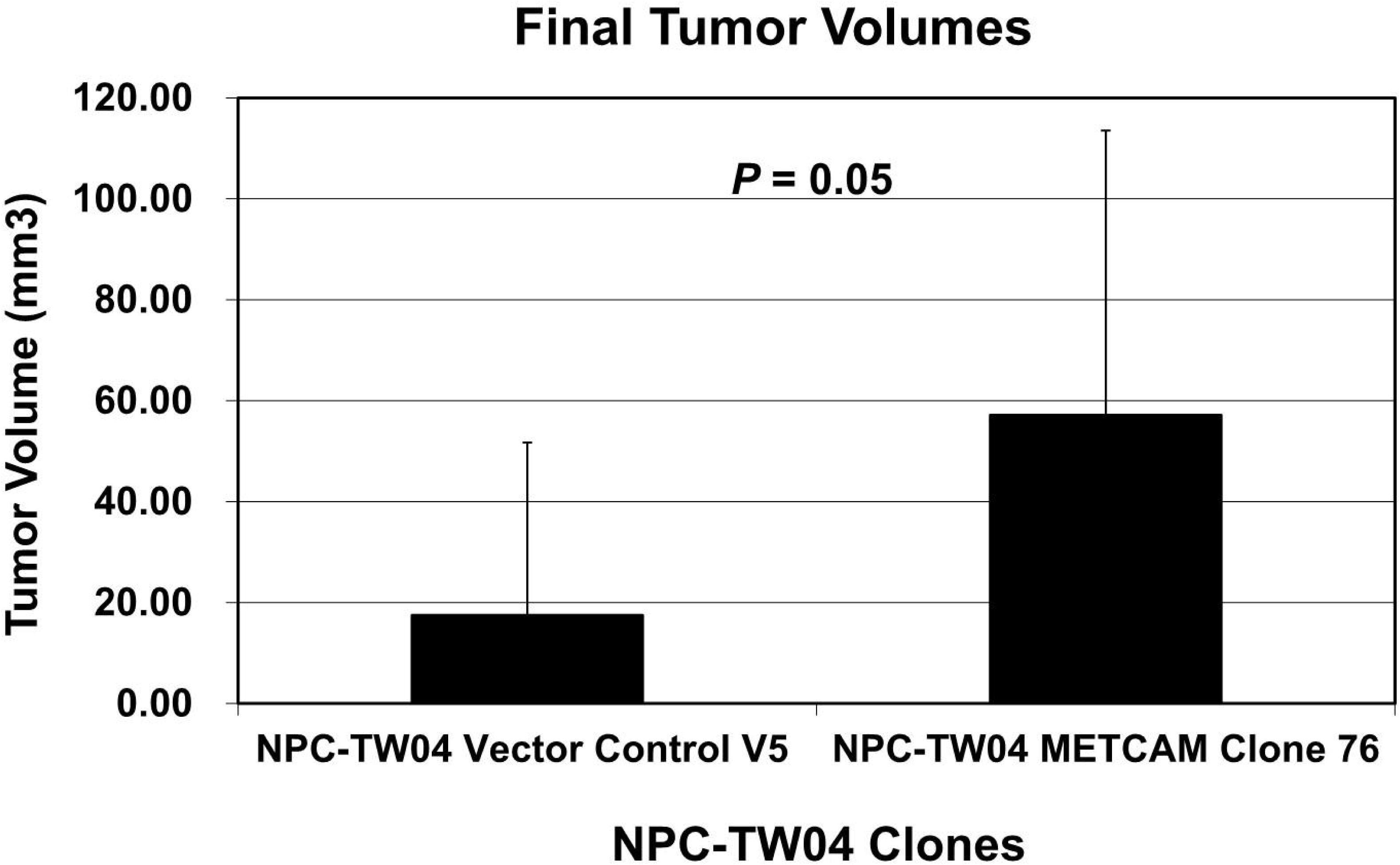
Effects of huMETCAM/MUC18 over-expression on the growth of the NPC-TW04 cells in the 3D basement membrane culture assay. The growth of the cells from the clone #76 and the control clone V5 in the 3D basement membrane assay was observed after 2-9 days and photographed with a SPOT digital camera attached to an inverted Nikon microscope. (A) shows the growth of the clone #76 and (B) shows the growth of the vector control V5.

### 3.5. Overexpression of huMETCAM/MUC18 promoted the *in vivo* tumorigenesis of the NPC-TW04 cells in athymic nude mice

To eliminate the possibility of yielding experimental artifacts by injection of a very high number of cells, about 10 x 10^7^ cells, which were used in the previous experiments [37, 38], we used a much lower cell number of cells for the injection. In the presence of Matrigel, injection of 5 x 10^5^ cells, tumor formation by the #76 clone was significantly more efficient than the V5 clone, as shown in Figure 5A. At the end point (40 days after injection) when the mice were terminated, the tumor-take and the final mean tumor volume of the clone #76 were 2.33 times higher and three times larger than that of the vector control clone V5, respectively, as shown in Figures 5B and 5C. From these results, we concluded that the over-expression of huMETCAM/MUC18 promoted the tumorigenicity and increased the final tumor volumes/weight of NPC-TW04 clones/cells in an athymic nude mouse model.

**Figure 5.**
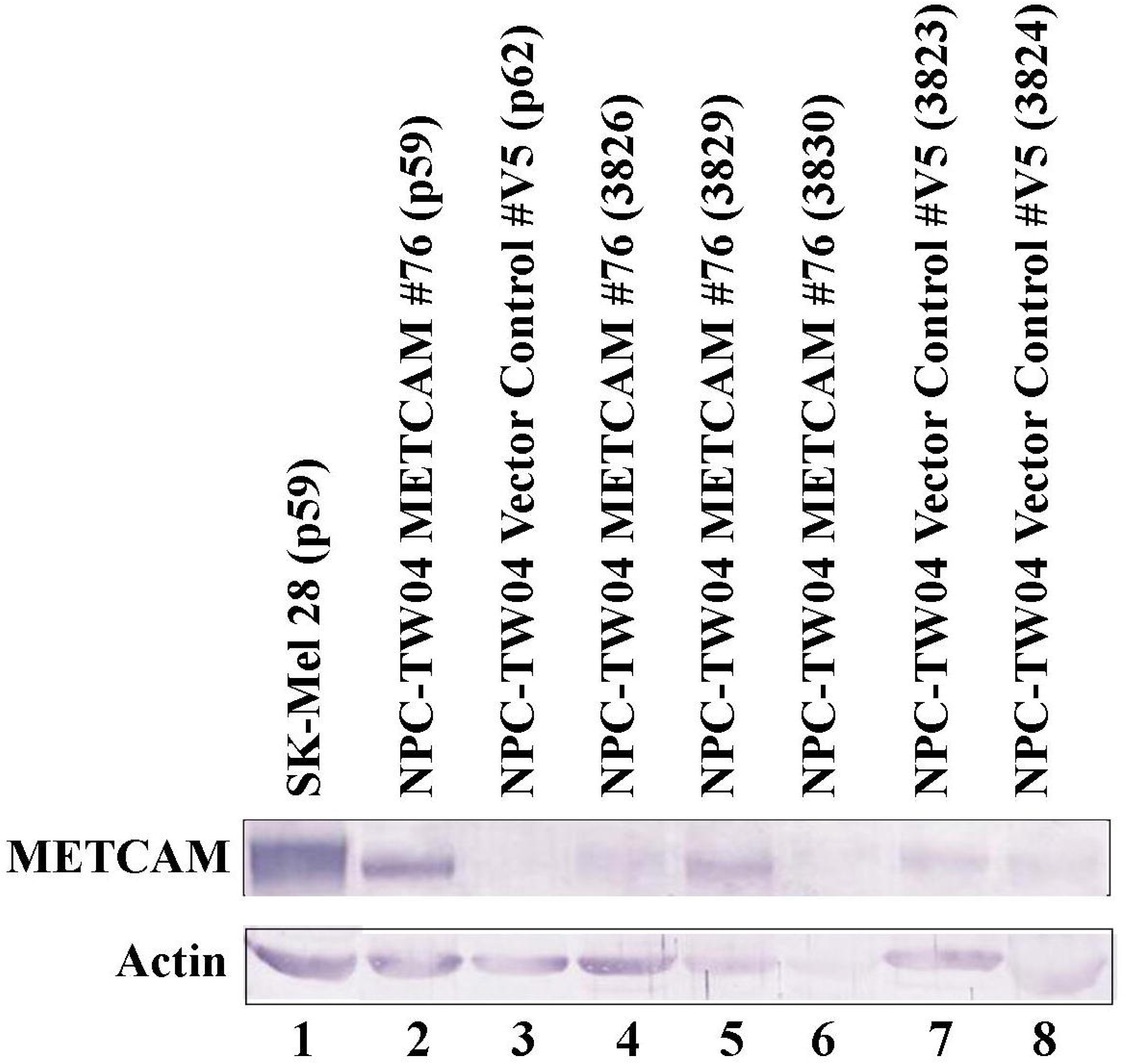
Effects of huMETCAM/MUC18 over-expression on the *in vivo* tumorigenesis of NPC-TW04 clones. Tumorigenicity of the METCAM clones #76 and the vector control clone V5 of NPC-TW04 cells was determined by subcutaneous injection of the cells from each clone in male athymic nude mice. (A) The tumor proliferation of the two clones by plotting mean tumor volumes versus time after injection of 5 x 10^5^ cells/mouse is shown. The tumor proliferation of the clone #76 was at least six times better than the vector control V5. *P* values were determined between tumor volumes through the time course of the METCAM clone #76 and that of the vector control clone V5. *P* values were 0.05 at all time-points, indicating that the differences between the clone #76 and the vector control clone V5 are statistically significant. (B) The tumor-take and (C) the mean final tumor volumes of the two clones in the athymic nude mice were compared at the endpoint. Both the tumor-take and the mean final tumor volumes/weights from five mice of the METCAM clone #76 were statistically significantly better than those of the vector control clone V5, respectively.

### 3.6. Expression of huMETCAM/MUC18 in tumor tissues

Figure 6A shows that the NPC-TW04 tumors derived from both clones #76 and V5 expressed different levels of METCAM/MUC18, but the electrophoretic mobility of METCAM/MUC18 expressed in the tumors was similar to that in the two clones grown *in vitro* in culture and that in the control cell line, SK-Mel-28. We concluded that the proteins expressed in tumors were not drastically modified or altered from the *in vitro* cultured clones/cell lines.

**Figure 6.**
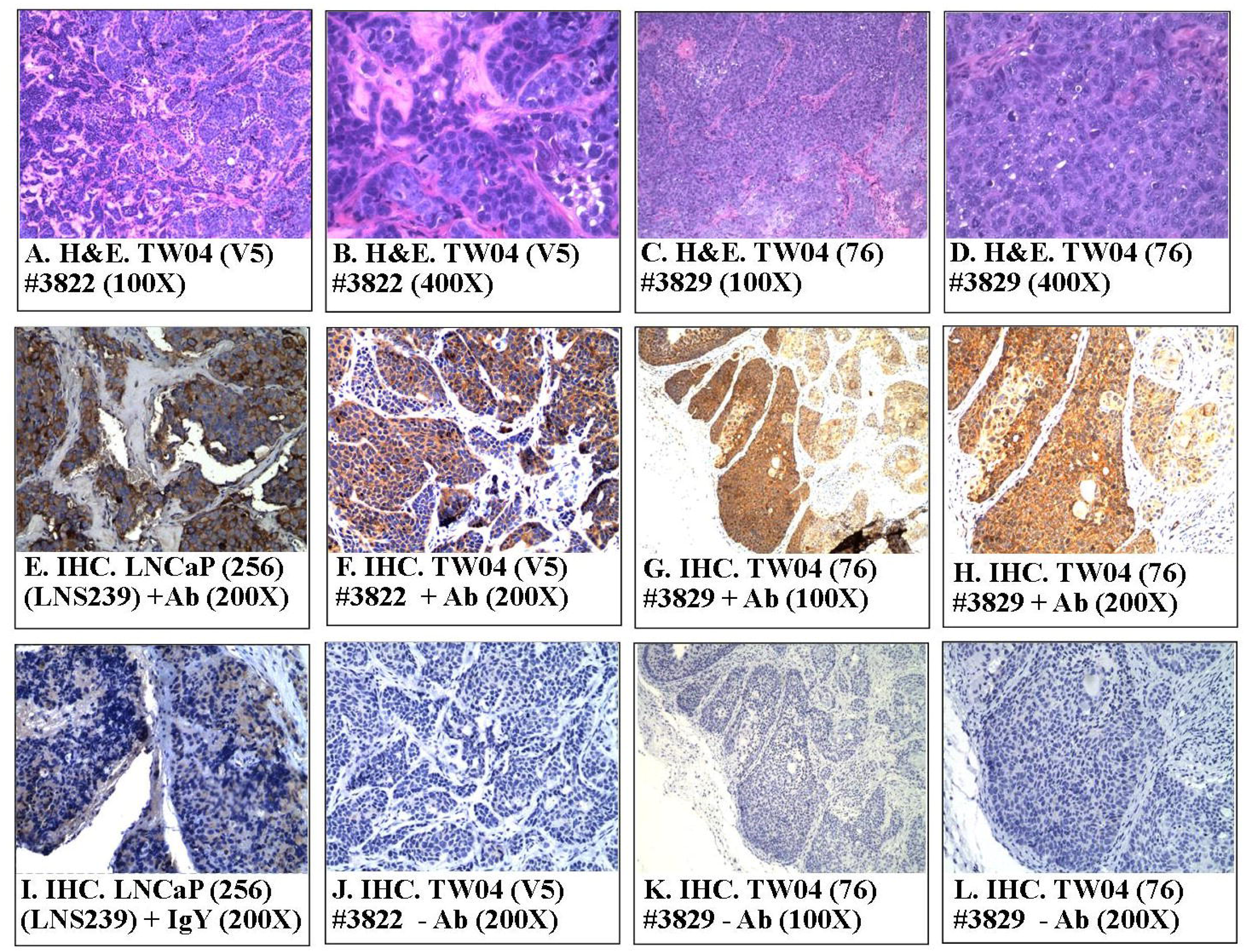
HuMETCAM/MUC18 antigen expression in tumor lysates and in tumor tissue sections. (A) The expression of huMETCAM/MUC18 in the lysates from the tumors was determined by Western blot analysis as described in the Section 2. The amount of protein loaded for each lane was 10 μg/lane. The expression of huMETCAM/MUC18 in the lysates from in vitro cultured SK-Mel-28 cells (lane 1) and from in vitro cultured NPC-TW04 clones #76 (lane 2) and V5 (lane 3) were used as the controls. The huMETCAM/MUC18 expression levels in the tumor lysates are shown in lanes 4-8. The huMETCAM/MUC18 expression levels in the lysates from the tumors of the METCAM clone #76 (lanes 4-6) and in the lysates from the tumors of the vector control clone V5 (lanes 7-8) are shown. The actin expression was used as the loading control for each lane. (B) Histology and immunohistochemistry (IHC) of the tumors of the NPC-TW04 clone #76 and the vector control V5 are shown. Panels A and B show the histology of the tumor sections from the vector control clone V5 and panels C and D that from the clone #76. Panels E to L show the IHC of the tumor sections. A tissue section of SC tumors derived from the human prostate cancer LNCaP-expressing clone (LNS239) was used as a positive external control for IHC staining (Panel E). Panels E to H show the anti-huMETCAM/MUC18 antibody staining of the cells in the tumor sections and panels I to L the negative controls without the antibody. The tumor section from the METCAM clone #76 showed strong brown color staining in IHC when the antibody was added (Panels G & H), however, the tumor section from the vector control clone V5 showed a weak background staining (Panel F). Panels I to L show the corresponding negative controls which show no staining in the adjacent sections when no anti-body or when the control chicken IgY was added.

Figure 6B shows the histology (panels A-D) and immunohistochemistry (panels E-L) of the NPC-TW04 tumors. Tumors from the vector control clone V5 appeared to be more confined (Panels A-B). In contrast, tumors from the clone #76 were more spread out in larger clusters (Panels C-D). METCAM/MUC18 antigens were weakly expressed in the centers of tumor clusters, when tumors were induced from the control clone V5 (Panel F). In contrast, METCAM/MUC18 antigens were strongly expressed in the entire tumor sections when tumors were induced from the clone #76 (Panels G-H). As also shown in Panels G-H, METCAM/MUC18 antigens were predominantly expressed on the cytoplasmic membrane, similar to the positive control tumor section from the LNCaP-induced tumors (Panel E). The results show that the tumors were indeed induced from the injected NPC-TW04 clones/cells.

### 3.7. Possible mechanisms of huMETCAM/MUC18-induced tumor promotion

The mechanism by which METCAM/MUC18 expression affects tumorigenesis of NPC cells has not been studied. By deducing knowledge which we learned from tumorigenesis of the NPC-TW01 cell line [11] and other tumor cell lines induced by METCAM/MUC18, METCAM/MUC18 may affect tumorigenesis by cross-talk with many signaling pathways that regulate apoptosis (Bcl2 and Bax), proliferation (PCNA), survival (AKT), glucose metabolism (LDH-A), and angiogenesis of tumor cells (VEGF) [15, 17, 18, 20, 23, 24, 26, 31, 32, 34, 36, 46]. We determined the expression of levels of Bcl2, Bax, LDH-A, PCNA, VEGF, and the ratio of phospho-AKT/AKT in tumor lysates. PCNA (proliferating cell nuclear antigen) is one of markers for cellular proliferation, because of its essential role in DNA replication and repairs and cell-cycle regulation [47, 48]. AKT, also known as protein kinase B, is a serine–threonine kinase downstream of the phosphatidylinositol 3-kinase (PI3K) signaling pathway, and has three Akt isoforms that may individually affect different cancer cells to varying degrees, suggesting specific targeting of different Akt isoforms for different types of cancer [49]. Functioning as a major effector protein in the PI3K pathway, AKT modulates normal cellular physiology like cell growth, survival, motility, proliferation, differentiation, metabolism, and cytoskeletal reorganization. AKT is also one of the most hyperactivated kinases in human cancers, influencing various biological phenomena that are directly involved in tumorigenesis [50]. Indeed, the PI3K/Akt signaling pathway is one of the most frequently altered signaling network in human cancer [49, 50]. The phosphatidylinositol 3-kinase (PI3K)/AKT/mammalian target of rapamycin (mTOR) network plays a key regulatory function in cell survival, proliferation, migration, metabolism, angiogenesis, and apoptosis [51]. Figure 7 shows that Bcl2, LDH-A, PCNA, VEGF, and ratio of phospho-AKT/AKT were increased in tumor lysates of the clone #76 when compared to those of the vector control clone #V5. In contrast, Bax was decreased in the tumor lysates of the clone #76 in comparison with those of the vector control clone #V5 (Figure 7).

**Figure 7.**
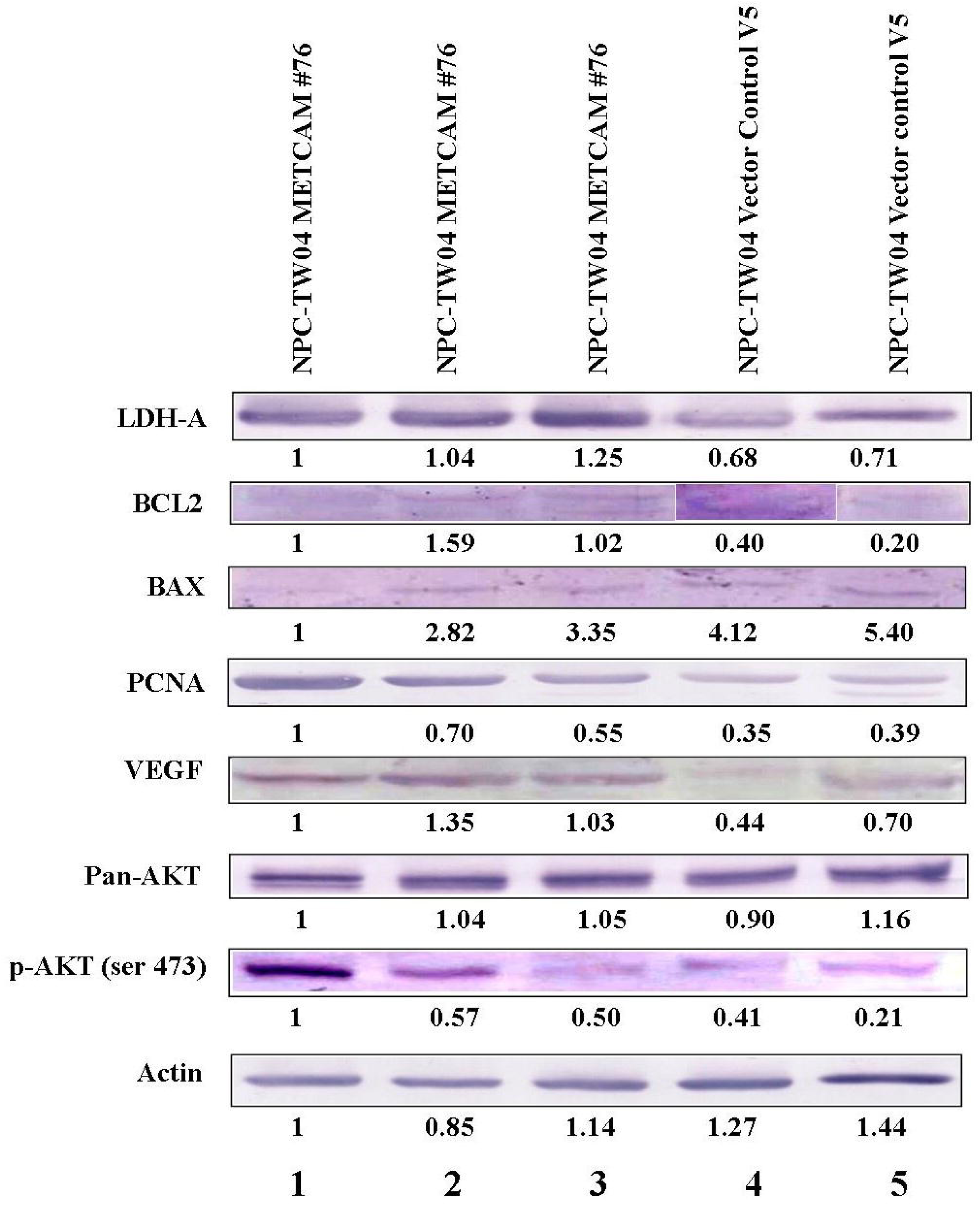
Effect of huMETCAM/MUC18 over-expression on the expression of key downstream effectors. Tumor lysates were used in the western blot analysis by using various antibodies, as described in the Section 2. (A) The Western blot results of levels of various key parameters, such as LDH-A, Bcl2, Bax, PCNA, VEGF, pan-AKT, and phospho-AKT(Ser473) are shown. (B) The quantitative results of the expression of the above effectors are shown.

Since VEGF level was elevated, as shown above, suggesting that angiogenesis was increased, the vascular density in the tumor sections was determined to further scrutinize this point. Figure 8 shows that the vascular density in the tumor sections induced from the NPC-TW04 clone #76, which overly expressed huMETCAM/MUC18, was indeed significantly higher than that of the vector control V5, further supporting that over-expression of huMETCAM/MUC18 may promote angiogenesis in the tumors.

**Figure 8.**
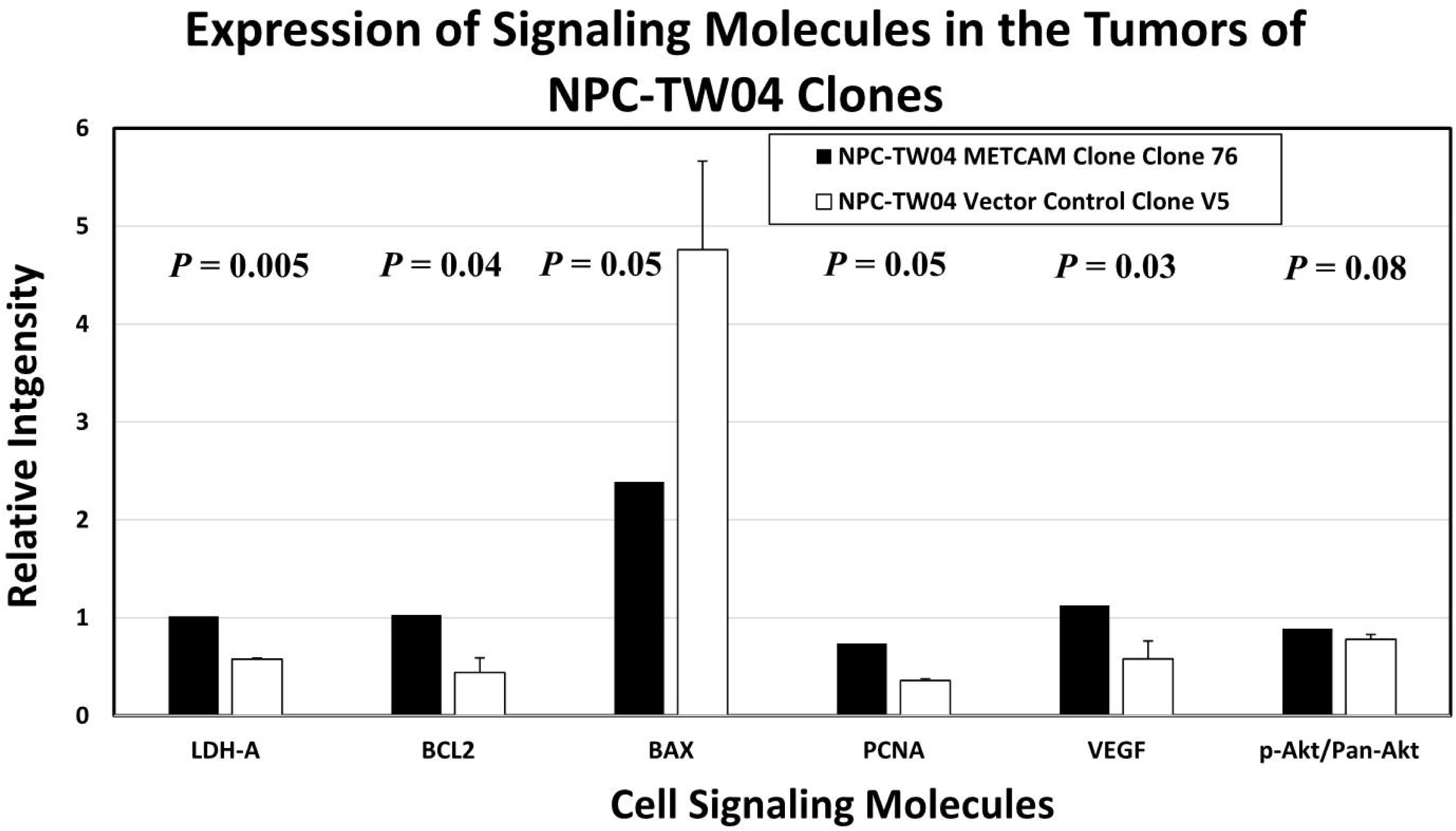
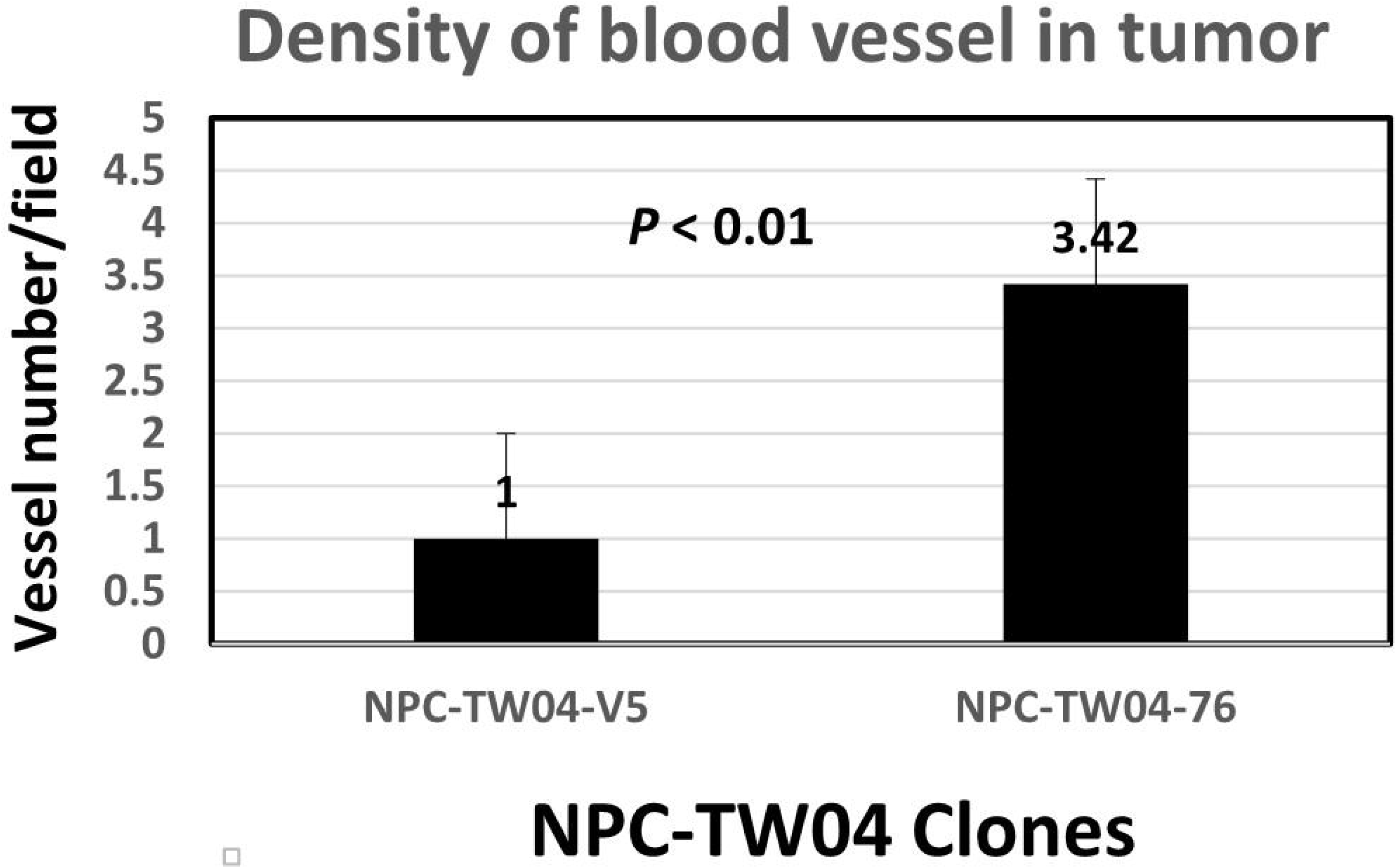
Vascular density of NPC-TW04 tumors. The vascular density was determined as described in the Section 2.

Taken together, we concluded that enforced expression of METCAM/MUC18 promoted the tumorigenesis of NPC-TW-04 cells by decreasing the pro-apoptotic index, Bax, and by increasing the indices of anti-apoptosis (BCL2 or the ratio of BCl2/Bax), aerobic glycolysis (LDH-A), proliferation (PCNA), the survival pathway (ratio of phospho-AKT/AKT), and angiogenesis (VEGF and vascularity density).

## 4. DISCUSSION

To further scrutinize the hypothesis deduced from the results of negative correlation of the expression of huMETCAM/MUC18 with the three histological types of NPC tissue sections [6], we tested effects of the over-expression of huMETCAM/MUC18 in NPC-TW04 cells, which were established from NPC type III, on their *in vitro* behaviors and *in vivo* tumorigenesis. We showed that the over-expression of huMETCAM/MUC18 increased *in vitro* motility and invasiveness (hence the epithelial-to-mesenchymal transition) and also increased the *in vitro* invasive growth in 3D basement membrane culture assay. We also showed that the over-expression of huMETCAM/MUC18 significantly promoted *in vivo* tumorigenicity of NPC-TW04 cell line. Since the huMETCAM/MUC18 in the tumor lysates was not drastically modified or altered, we suggest that the native (unaltered) form of huMETCAM/MUC18 was responsible for these effects. To understand the possible mechanism, we showed that the expression levels of downstream effectors, such as an anti-apoptosis index, a proliferation index, an index for a metabolic switch to aerobic glycolysis, an angiogenesis index and the survival pathway index were increased, whereas a pro-apoptotic index (Bax) was decreased, suggesting that huMETCAM/MUC18 may mediate tumor promotion by elevating anti-apoptosis, angiogenesis, proliferation, survival pathway, and metabolic switch to aerobic glycolysis and by reducing apoptosis. Taken together, we concluded that over-expression of huMETCAM/MUC18 in NPC-TW04 cells promoted the tumor formation induced by the cells, supporting the notion that huMETCAM/MUC18 plays a tumor promoter role in the development of the type III NPC [37, 38]. Therefore, the list of cancers in which METCAM/MUC18 plays a tumor promoter role has been extended from breast cancer, prostate cancer, osteosarcoma, and angiosarcoma [46] to NPC type III (this report).

In contrast to the previous conclusion drawn from the results of various *in vitro* and *in vivo* tests of the NPC-TW01 cells, which were established from the NPC type I [11], surprisingly in this report the conclusion drawn from the results of similar tests of the NPC-TW04 was completely opposite to that of the NPC-TW01 cells in that over-expression of METCAM/MUC18 increased the development of NPC type III. This suggests that the negative correlation in the incidence of NPC type III was fortuitous [10]. Interestingly, this is another example of confounding the true role of METCAM/MUC18 in the progression of a human cancer in that the conclusion or hypothesis drawn from a positive correlation of clinical prognosis with the increased expression of METCAM/MUC18 in malignant ovarian cancer specimens was wrong without the support of evidence from more extensive *in vitro* tests and *in vivo* tests in an animal model [32–35]. Taken together, this points to an important concept that a conclusion or hypothesis derived from the correlation with pathology should be further scrutinized by direct evidence obtained from extensive tests of *in vitro* growth behavior and *in vivo* tests in an animal model system [34].

The dual role of huMETCAM/MUC18 in the progression of NPC appears to be consistent with the notion in that the dual role of huMETCAM/MUC18 in the progression of human cancer is generally found in different cell lines of the same cancer type or different cancer types, suggesting that huMETCAM/MUC18 affects the tumorigenesis and metastasis of epithelial tumor cell lines in a very complex way. We did not know the exact reasons, but we suggest that one of which may be due to different co-factors present in different cancer cell lines, which may modulate the role of huMETCAM/MUC18 in these processes via homophilic interactions with endothelial cells and immune cells and via heterophilic interactions with other cells and also with the extra-cellular matrix in the tumor microenvironment [46], though the corresponding heterophilic ligands of huMETCAM/MUC18 in these cancer cell lines are yet to be identified [46]. The other likely mechanisms, such as mis-sense mutation, different post-translational modifications, and altered subcellular localization [52], may not be excluded and thus should also be investigated in the future.

One point worth of noting is that the number of tumor cells used for injection into animal model appears to be critical for revealing the true effect of the overly expressed METCAM/MUC18 on tumorigenesis, since artifacts may result from injecting too high a number of tumor cells [44]. At the smaller cell number, the difference in the tumorigenicity of the high-expression clone versus the vector control was clearly shown in this report. Since the interactions of tumor cells with the heterogeneous and complex components in the tumor microenvironment significantly contribute to the development of cancer [53, 54]. Initial number of cancer cells injected could affect the interactions of the tumor cells with the surrounding immune cells, tumor-associated fibroblasts, blood vessels, extracellular matrix (ECM), lymphocytes, bone marrow-derived inflammatory cells, and signaling molecules in the tumor microenvironment and the manifestation of their stemness. Especially since METCAM/MUC18 is an integral membrane protein with the cell adhesion function [55] as well as it is also expressed in mesenchymal cells [56], its over-expression in these NPC cells should have a profound effect on how tumor cells interact with the various cells in the tumor microenvironment. As such, a small number of tumor cells injected may be more sensitive to the delicately regulated physiological conditions in the tumor microenvironment and reflect the true initial tumorigenesis, whereas a large number of the tumor cells may secrete over-dosed small molecules to overwhelming influence the over-powering dynamic interactions of the initial tumor cells with those cells in the tumor microenvironment in an artificial and non-physiological manner and thus, mask the dramatic effect of METCAM/MUC18 on the tumorigenesis of these NPC cells [53, 54].

## 5. CONCLUSIONS

HuMETCAM/MUC18 was clearly proven in this report to play a tumor promoter role in the progression of the NPC-TW04 cells, which were established from the NPC type III. The tumor promoting effect of huMETCAM/MUC18 on these NPC cells was not due to increasing the intrinsic *in vitro* growth rate of the NPC-TW04 cells, but it rather might be due to the augmentation of the intrinsic *in vivo* growth via decreased apoptosis and concomitantly elevated anti-apoptosis, proliferation, survival pathway, rewiring metabolic switch from oxidative phosphorylation to aerobic glycolysis, and angiogenesis. The knowledge learned from the studies in this report and the parallel studies in the published report [11] should be useful for broadening our understanding the progression of different types of NPC; furthermore, they may also be useful for designing different therapeutic means for the control and the clinical treatment of different types of NPC. Four general strategies may be used for the designing therapeutic means to treat the NPC type III patients: (a) immunotherapy by using anti-METCAM/MUC18 monoclonal antibodies [28, 57], (b) treatments by using METCAM/MUC18-specifc siRNAs [27, 28, 58], (c) blocking the modulating role of the specific heterophilic ligand(s) [46], and (d) blocking the key members of the downstream pathways that are activated by this tumor promoter [46].

## Abbreviations

3D: three-dimensional;
CAM: cell adhesion molecule;
EBV: Epstein-Barr virus;
huMETCAM: human METCAM;
Ig: immunoglobulin;
IHC: immunohistochemistry;
IP: intraperitoneal;
METCAM: metastasis regulation cell adhesion molecule;
NPC: nasopharyngeal carcinoma.

## Author Contributions

GJW and YJC conceived and designed the research; YCL and CCK performed the research, acquired the data and analyzed and interpreted the data. All authors were involved in drafting and revising the manuscript. All authors have read and agreed to the published version of the manuscript.

## Institutional Review Board Statement

All animal studies were conducted in accordance with the Declaration of Helsinki, and approved by approved by the Mackay Memorial Hospital Experimental Animal Care and Use Committee (an approval number MMH-A-S-101-31, 2013) and Chung Yuan Christian University Experimental Animal Care Committee (an approval number10119, 2013).

## Informed Consent Statement

Not applicable

## Data Availability Statement

The datasets used and/or analyzed during the current study avail-able from http://www.lib.cycu.edu.tw/thesis with the consent from the corresponding author upon reasonable request.

## Acknowledgments

We thank Mr. Jie-Hong Chen for helping us using One-Way ANOVA for statistical analyses and Mr. Jonathan Wu and Mrs. Mei Wu for critical proof-reading English.

